# *Stentor stipatus*, a new unicellular species, demonstrates novel phototaxis and habituation

**DOI:** 10.1101/2024.08.03.606273

**Authors:** Deepa H. Rajan, Benjamin Lee, Ashley Albright, Eric Tang, Kristen Ressler, Arnie Maravillas, Carmen Vargas, Wallace F. Marshall, Daniel B. Cortes

## Abstract

**Background:** *Stentor*, the genus of large trumpet-shaped ciliates, is well-known for its complex morphology and striking behaviors. Members of this genus are distributed throughout the world in a wide and diverse pool of freshwater ecosystems. Recently, the molecular phylogeny of *Stentor* has been explored through comparison of 18S small subunit (SSU) ribosomal DNA (rDNA) sequences, clarifying several previously mischaracterized species and species complexes. However, despite their wide distribution, to-date, only about a dozen species of *Stentor* have been described and verified by phylogenetic means.

**Results:** Here, we introduce the discovery of a new species within genus *Stentor*: *Stentor stipatus* spec. nov., so named for their distinctive cytosolic dark pigmented granules which surround the macronucleus and are also present cortically alongside cortically-distributed green microalgae. We present morphological, phylogenetic, ecological, and behavioral characterizations of these cells. Phylogenetic analysis of *S. stipatus* spec. nov. by comparison of SSU rDNA sequence suggests it is a distinct species from its closest relative, *S. amethystinus*. We demonstrate that *S. stipatus* spec. nov. is capable of habituation in response to repeated mechanical stimulation. Further, *S. stipatus* spec. nov. exhibits strongly directed positive phototaxis, like its relative S. pyriformis, but with a distinct action spectrum from both *S. coeruleus* and *S. pyriformis*. Finally, *S. stipatus* phototaxis response strength varies in a consistent pattern throughout the day, providing evidence of potential circadian regulation.

**Conclusions:** This work expands the current understanding of the ecological distribution of and behavioral features present within genus *Stentor*.

## BACKGROUND

Ciliates (phylum Ciliophora) (Lee and Kugrens, 1992) are a major component, by biomass, of most aquatic ecosystems (Lischke *et al*., 2016). Ciliates are effective predators of smaller microorganisms and also serve as an important food source for metazoans, providing a significant link between prokaryotic and eukaryotic food chains (Porter *et al*., 1979; Liu *et al*., 2016). Many ciliates also contain photosynthetic endosymbionts while retaining their ability to ingest prey organisms, creating further links in the metabolic chain. Among the ciliates, the genus *Stentor* is known for its trumpet-shaped morphology and large size (up to several millimeters in length), with some species visible to the naked eye (Tartar, 1961; Slabodnick *et al*., 2014). The name *Stentor* references the Greek herald in the *Iliad* (Slabodnick *et al*., 2014) with a “voice as powerful as fifty voices of other men” (Homer), alluding to the cell’s trumpet shape. *Stentor*, like other members of the Heterotrichae class, are highly contractile and have a prominent membranellar band comprising long cilia around the oral cavity as well as shorter cilia distributed over the rest of the body (Tartar 1961).

Historically, *Stentor* species were described through characterization of several key morphological traits; (1) the presence or absence of endosymbiotic microalgae, (2) the shape of the macronucleus, (3) the presence of and color of pigment granules in the cortex, and in some instances (4) the number and order of somatic kineties. There are around a dozen known species of *Stentor*, and all are characterized by trumpet-shaped morphology (Foissner and Wölfl, 1994), and verified by phylogenetic analysis using alignment of the 18S small subunit (SSU) ribosomal DNA (rDNA) sequence (Thamm *et al*., 2010; Fernandes *et al*., 2016).

*Stentor* can vary significantly in their morphologies. Some species of *Stentor*, like *S. amethystinus*, are shorter and fatter, while others like S. polymorphus have a more elongated shape (Foissner and Wölfl, 1994; Foissner and Berger, 1996). Several species are heavily pigmented like *S. niger*, while others like *S. muelleri* and *S. roeselii* are transparent and seemingly devoid of pigment granules (Foissner and Wölfl, 1994). A few species, such as *S. roeselii* and S. muelleri, may build lorica, which are mucilaginous structures surrounding the base of the cell into which the *Stentor* can contract (Foissner and Wölfl, 1994). Other species, like *S. pyriformis*, are known to have algal endosymbionts (Foissner and Wölfl, 1994; Hoshina *et al*., 2021; Boudreau *et al*., 2025). However, most ecological work on the biogeographic distribution of *Stentor* has focused on *S. coeruleus* and *S. pyriformis*, while other members of the genus remain relatively understudied. Overall, little is known about the natural history of *Stentor* as a whole, or about the evolutionary relationships among the various *Stentor* species, or about the ecological distribution of these species.

*Stentor* have a rich history as a model of unicell behavioral research; *S. coeruleus*, amongst other species, have been studies extensively for their ability to perform complex behaviors, such as phototaxis (Tartar, 1961; Wood, 1976; Kim *et al*., 1984), decision-making (Dexter *et al*., 2019), and learning (Rajan *et al*., 2023b) despite being unicellular, and thus lacking a nervous system. Furthermore, *Stentor* have the remarkable ability to regenerate after being injured by microsurgery (Morgan, 1901; Tartar, 1961; Marshall, 2021). Studying *Stentor* thereby has the potential to reveal fundamental insights about cellular computation, wound healing, and the evolution of complex morphology and cellular behaviors.

A notable form of computation that *Stentor* cells exhibit is habituation, which is a form of learning characterized by a decrease in response after repeated stimulation. Single cell habituation has primarily been studied in *S. coeruleus*, which contract in response to mechanical stimulation, as an apparent escape response from predators in their natural pond habitat. However, after repeated stimulation, *S. coeruleus* cells habituate and thus stop contracting (Rajan *et al*., 2023b). The mechanism by which habituation occurs is not yet known, and investigating this process in a set of related species is one way to get new clues about the process and its components.

Here, we introduce the discovery of a new species within this genus: *Stentor stipatus* spec. nov., so named for their distinctive dark pigment granule aggregates. We present morphological, phylogenetic, ecological, and behavioral characterizations of this new *Stentor* species. These findings expand our understanding of *Stentor* species diversity, both in terms of their ecology and behavioral complexity, and highlight the importance of continued exploration to uncover hidden microbial biodiversity awaiting discovery in even well-studied ecosystems.

## METHODS

### Collection and culture of the organisms

*Stentor stipatus* spec. nov. samples were collected from the white cedar swamp within the Two Ponds Conservation Area, near the edge of Jones Pond in Falmouth, Massachusetts (approximate coordinates of 41°33’52.0”N 70°36’26.9”W) continuously between early May and mid-July of 2023 and 2024. Collection involved disturbing the underwater topsoil and scooping a portion of the suspended sediment and plant matter into small wide-mouth collection jars. Due to their heavy dark coloration, *S. stipatus* spec. nov. cells were readily identifiable by the naked eye and through use of a low-magnification SM2-TZ stereomicroscope (AmScope) with transillumination. *S. stipatus* spec. nov. cells were concentrated by a bright light source and physically isolated from collection samples through use of a pipette. Isolated cells were then placed in either pasteurized water collected from the same source, or synthetic pond water (SPW) made to approximate the chemical composition of the source water.

Initially, *S. stipatus* spec. nov. cultures were grown in 100 ml wide-mouth jars at room temperature on a windowsill that received around 10-12 hours of natural sunlight each day. Cultures were fed approximately 0.5 ml of *Chilomonas sp*. culture (Carolina Biologicals Cat. 131734) twice weekly. Under these conditions, *S. stipatus* spec. nov. cultures remained stable for approximately three months, while at the Marine Biological Laboratory in Woods Hole, Massachusetts.

Long-term culturing of *S. stipatus* spec. nov. was achieved by growing cultures in SPW. Cultures were kept in a low temperature incubator, at 18 °C under 12 hours of direct white light (∼5000K), from 7AM to 7PM, and 12 hours of darkness from 7PM to 7AM (hereafter referred to as 12L/12D conditions). *S. stipatus* spec. nov. cultures were fed *Chilomonas* as described above, and have been maintained consistently in the lab for over 18 months in this manner.

*S. coeruleus* cells used for habituation experiments were cultured using the methods described in Rajan *et al*. 2024. Cells were fed *Chlamydomonas* and were grown in pasteurized spring water, (Carolina Biological 132458; Burlington, NC), which contains 8.6 mg/l Cl, 13.89 mg/l Na, 4.15 mg/l Mg, 0.12 mg/l S, 15.51 mg/l Ca, 2.50 mg/l K, 0.09 mg/l Sr, 0.06 mg/l F, 20196 µg/l Si, 53.21 µg/l Ba, 1.14 µg/l Cr, 1.72 µg/l V, 9.28 mg/l NO3, 20.59 µg/l P, 0.06 mg/l PO4 (ICP-OES analysis; ATI-Aquaristik) (Table 1).

**Table 1.** Chemical composition of water from several different sources used in this study. Listed in concentration (mg/L) based on measures taken from ICP-OES analysis. WCS = white cedar swamp, Jones = Jones Pond, one of the ponds bounding the white cedar swamp of Falmouth, Pandapas = Pandapas Pond in Blacksburg, VA – this water was used for initial culturing attempts of *S. stipatus* spec. nov. PSW = Carolina Biological pasteurized spring water, commercial water used for culturing of habituating *S. stipatus* spec. nov. and *S. coeruleus*.

| Element | WCS | Jones | Pandapas | PSW |
| --- | --- | --- | --- | --- |
| Cl | 12.705 | 17.24 | 11.25 | 8.6 |
| Na | 14.37 | 16.81 | 13.33 | 13.89 |
| Mg | 0.42 | 2.51 | 1.88 | 4.15 |
| S | 0.135 | 1.98 | 0.55 | 0.12 |
| Ca | 4.94 | 3.51 | 3 | 15.51 |
| K | 4.095 | 3.82 | 2.89 | 2.5 |
| Sr | 0.045 | 0.02 | 0.01 | 0.09 |
| B | 0.01 | NA | NA | NA |
| F | 0.525 | 0.61 | 0.18 | 0.06 |
| Si | 2.648 | 4.266 | 2.565 | 20.196 |
| Ba | 0.010095 | 0.01728 | 0.02559 | 0.05321 |
| Ni | 0.00111 | NA | NA | NA |
| Mn | 0.01109 | NA | NA | NA |
| Cr | 0.00661 | NA | NA | 0.00114 |
| Fe | 1.8605 | NA | 0.08116 | NA |
| Cu | 0.003165 | 0.00206 | 0.00373 | NA |
| Ag | 0.010475 | NA | NA | NA |
| V | 0.0012 | NA | NA | 0.00172 |
| Zn | 0.013585 | 0.0045 | 0.00293 | NA |
| NO<sub>3</sub> | 1.62 | 0.6 | NA | 9.28 |
| P | 0.050155 | NA | NA | 0.02059 |
| PO<sub>4</sub> | 0.137 | NA | NA | 0.06 |
| Al | 0.05563 | NA | 0.01549 | NA |

### Chilomonas culturing

*Chilomonas sp*. was initially grown in 100 ml wide-mouth jars in pasteurized pond water collected from a local pond (Morse Pond in Falmouth, Massachusetts) during the 2023 season. Cultures from 2024 onwards were maintained with the same SPW used to culture *S. stipatus* spec. nov. Feed cultures were seeded with 2-3 boiled wheat berries to provide nourishment. Cultures were kept in the dark at 20 °C and were checked regularly for any potential contaminants such as rotifers or other ciliates; contaminated cultures were disposed of.

### Preparation of Synthetic Pond Water (SPW)

Three separate samples, each containing triplicate 15ml tubes full of pond water from the white cedar swamp of Falmouth, MA, were sent for water composition analysis through ICP-OES (ATI-Aquaristik; Table 1). Major chemicals (those above 10 µg/L) were averaged across all three samples to generate an average chemical composition. Using these concentrations, we put together a list of salts and fertilizers that could be combined to create the same overall composition. These components (Table 2) were mixed into a 10X concentration with milliQ lab water. A 1X solution of SPW was prepared fresh as needed by diluting 10X SPW 10-fold in milliQ water.

**Table 2.** Chemical composition of synthetic culturing media listed in mM concentrations as well as mg/L and 10X mg/L, based on average measurements taken from ICP-OES analysis of the white cedar swamp.

| Chemical | mM | mg/L | 10x mg/L |
| --- | --- | --- | --- |
| NaCl | 0.367 | 21.44748 | 214.4748 |
| NaNO3 | 0.026 | 2.20986 | 22.0986 |
| K2HPO4 | 0.0017 | 0.29614 | 2.9614 |
| KOH | 0.098937 | 5.55091 | 55.5091 |
| NaF | 0.026316 | 1.10495 | 11.0495 |
| NaOH | 0.143307 | 5.73186 | 57.3186 |
| CaO3Si | 0.1 | 11.616 | 116.16 |
| Na2O3S | 0.004367 | 0.55044 | 5.5044 |
| FeSO4 | 0.03402 | 9.45782 | 94.5782 |

### Morphological observations

General shape and size descriptions were made using data acquired on a Zeiss AxioZoom V.16 equipped with a PlanNeoFluar Z 2.3x/0.57 objective and a Thorlabs CS505CU 12-bit color CMOS camera (Thorlabs) and an Amscope stereomicroscope (AmScope SM2-TZ) with a Thorlabs monochrome CMOS (CS135MUN; Thorlabs). Images and videos were acquired using ThorCam software (Thorlabs). The scale for size measurements was established using a micrometer slide (AmScope) which was imaged at maximum magnification with each microscope.

### Fixation and fluorescence imaging of *S. stipatus* spec. nov

*S. stipatus* spec. nov. cells were collected from sample cultures and washed in fresh SPW. 10-20 cells were then moved into ∼100 µl of SPW onto a #1.5 22×2mm coverslip which was covered with a 1mm thick silicone pad with a 10mm circular hole cut out of it. The silicone pad with a circular hole created a spacer such that *Stentor*s placed in the central circle cutout could be sandwiched between coverslip and glass slide without any compression. After cells were moved into the sample area of the silicone-padded coverslip, 100 µl of relaxation buffer [10 mM EGTA, 70 mM Tris, 3 mM MgSO4, 7.5 mM NH4Cl, 10 mM phosphate buffer (pH 7.1), pH to 7.2] was added and mixed. Cells were incubated for 10-15 minutes in relaxation buffer, until all cells were visibly extended. Following relaxation, most of the liquid was removed by pipette and 200 µl of 1% glutaraldehyde in PBS was added to the cells. After 30-60 minutes in glutaraldehyde, cells were washed twice, for 5 minutes each, with 200 µl of PBS. Next, PBS was removed and replaced with 200 µl of PBST (Tween-20). After 5 minutes in PBST, most of the liquid was removed from and replaced with ∼50 µl of Prolong Gold + DAPI (Thermo Fisher) mounting solution. After the addition of Prolong Gold, cells were maneuvered using an eyelash tool to ensure no clumping and that all cells were settled at the bottom of the liquid volume close to the coverslip surface. A slide was then inverted onto the coverslip, taking care not to trap any bubbles within the sample area. Excess liquid was wicked away and the slide was left inverted overnight in a 4 °C refrigerator. After overnight drying, the slide was turned right side up briefly and sealed with nail polish, then re-inverted and stored coverslip down at 4 °C indefinitely.

Imaging was carried out on a Nikon Eclipse T2 inverted microscope body with an Illumina 7-laser module (Illumina) and an X-Light V3 spinning disk module (CrestOptics) with a 100x silicone immersion objective and a Prime 95B sCMOS camera (Teledyne Photometrics) using Nikon Elements software. Algal cells within the *Stentor* cells were autofluorescent most strongly at 561 nm and 640 nm. The *S. stipatus* spec. nov. cells themselves were autofluorescent at 365 nm, 440 nm, 488 nm, 514 nm, 561 nm, and 640 nm.

### DNA extraction and amplification

Single cells were washed 3 times in pasteurized spring water (Carolina Biologicals Cat. 132458). DNA was isolated from these cells using a combination of worm lysis buffer (50 mM KCl, 10 mM Tris pH 8.2, 2.5 mM MgCl_2_, 0.45% Igepal, 0.45% Tween 20, 0.01% gelatin) and proteinase K (New England Biologicals P8107) at a ratio of 166:1. Each *Stentor* cell was individually transferred in minimal volume into a tube containing 2.5 µL the worm lysis buffer + proteinase K. The tube was then placed in a -80 °C freezer for 10 minutes and subsequently kept on ice. To complete the lysis, the samples were then heated at 60 °C for 60 minutes and then at 95 °C for 15 minutes.

For the polymerase chain reaction (PCR), a master mix solution was prepared containing 75 µL of Q5 High-Fidelity 2X Master Mix (New England Biologicals M0492S), 7.5 µL of the forward primer (5’-CAGCAGCCGCGGTAATTCC-3’), 7.5 µL of the reverse primer (5’-CCCGTGTTGAGTCAAATTAAGC-3’), and 45 µL of nuclease-free water (New England Biologicals B1500S). 22.5 µL of the master mix solution was then pipetted into each tube containing ∼2.5 µl of purified clonal culture DNA. PCR cycling parameters followed the conditions described in New England Biolabs 2022. Single band PCR products were confirmed using agarose gel electrophoresis, cleaned and concentrated (Zymo Research D4066), and then sent for Sanger sequencing.

### Phylogenetic analysis

The 18S SSU rDNA sequence was selected for phylogenetic comparison of *S. stipatus* spec. nov. to other *Stentor* based on its wide use as a phylogenetic marker for *Stentor* and other ciliate phylogenies (Thamm *et al*., 2010; Fernandes *et al*., 2016; Hoshina *et al*., 2021). Published 18S SSU sequences previously used for assembly of *Stentor* phylogenies were identified by their accession IDs (Thamm *et al*., 2010; Fernandes *et al*., 2016; Taher *et al*., 2020) and downloaded in FASTA format from GenBank. We also included sequences from *Maristentor* dinoferus, *Blepharisma americanum, Blepharisma hyalinum*, and *Condylostentor auriculatus* as outgroups. We generated four 18S SSU rDNA sequences from *S. stipatus* cells (PX056132, PX056133, PX056134 and PX393983), along with single sequences generated from wild isolates of *S. coeruleus* (PX056129), *S. muelleri* (PX056130), and *S. pyriformis* (PX056131), all non-*S. stipatus* spec. nov. cells were cultured as described before (Lin *et al*., 2018; Boudreau *et al*., 2025). DNA isolation and sequencing for these was carried out as described in the “DNA extractionand amplification” methods section. Sequence files were trimmed with ChromasPro (v2.2.0; Technelysium Pty. Ltd.) using default trimming options to improve read quality, and aligned using MAFFT (version 7) under default settings. Alignments were used in IQTREE (multicore version 2.0.7) to infer a maximum-likelihood (ML) tree that was rooted using C. auriculatus as an outgroup. The ‘automatic’ substitution model method was used to identify the optimal nucleotide substitution model, TN+F+I+G4. Branch reliability was assessed using bootstrapping with 10,000 replicates as well as 1000 replicates of single-branch Shimodaira-Hasegawa approximate likelihood ratio test (SH-aLRT) (Guindon *et al*., 2010), and single-branch Bayesian-like transformation of aLRT (aBayes(Anisimova *et al*., 2011).

### Habituation experiment

Habituation was assessed in the *Stentor cells* using a modified version of the protocol described previously (Rajan *et al*., 2023a; Rajan and Marshall, 2025). We used an Arduino-controlled habituation device to deliver mechanical taps to the cells at a specified force and frequency. We took videos of the cells over the course of the habituation experiment and quantified the fraction of cells contracted. We only quantified anchored *S. stipatus* cells with a visible posterior region, i.e., cells that were angled to reveal their slightly oblong shape, and excluded cells that were positioned such that only a top-down spherical view was visible from the microscope. This was because contractile responses are only possible to ascertain when the posterior region is visible to the microscope. The change from a slightly oblong cell shape to a spherical shape immediately after delivery of a mechanical tap was counted as a single contraction. We also excluded cells that detached from the bottom of the petri dish after receiving a mechanical tap because this detachment is unlikely to be a behavioral response but rather a dislodgment caused by the shearing force produced by the mechanical perturbation.

A reduction in the proportion of cells that contracted over time indicated successful habituation, as the cells learned to ignore the mechanical stimulation. Contraction and habituation in *S. coeruleus* are stimulus specific, such that cells that habituate to mechanical stimuli can still respond to electrical and photic stimuli (Wood, 1973). This suggests that observed reductions in contraction responses in *Stentor* species likely result from habituation to specific stimuli rather than from fatigue. Furthermore, as described in Rajan *et al*. 2023a, we verified that contractions occurred primarily immediately after the delivery of a mechanical tap and are thus very unlikely to be spontaneous contractions due to external stimuli.

To improve video resolution, instead of using a USB microscope, the habituation device was placed directly on the stage of a dissection microscope (Zeiss Stemi 305) and an iPhone camera was affixed directly to the microscope eyepiece. The 35 mm petri dish on the habituation device contained 26 lab-grown *Stentor* coeruleus (VWR 470176-586) cells and 100 wild-caught *Stentor stipatus* spec. nov. cells. The first two replicates for the *Stentor* coeruleus habituation experiments were carried out with cells from the same culture, and the final replicate was conducted with cells from a different culture. After two hours of acclimation, the cells were trained with level 3 pulses (higher force) at a frequency of 1 tap/min. Force levels are defined in Rajan *et al*. 2023a: level 1 is the highest force and level 5 is the lowest force. Level 3 taps, employed in these experiments, result in the downward displacement of ∼2 mm and a downward peak force of ∼0.557 N (Rajan *et al*., 2023a).

### Phototaxis and action spectrum assays

Phototaxis of *S. stipatus* spec. nov. was assessed using a modified version of the protocols previously employed for other *Stentor* species (Kim *et al*., 1984). A custom 3D-printed chamber (Figure S1A) was made using high-clarity transparent resin (SirayaTech Blu V2). Following initial printing, the chamber was washed in 99% isopropanol for 5 minutes and then exposed to strong UV light for 30 minutes to remove residual resin and fully harden the structure. Following the long cure, the chamber was incubated at 60 °C for 24 hours and soaked in pasteurized pond water overnight to remove any remaining volatile chemicals which may be detrimental to *Stentor* health. The chamber design featured a space under the specimen area, into which a high-powered LED (Chanzon) of specific wavelength could be inserted. Along the length of the specimen area, clear demarcation lines were added that split the long column into quadrants for easier quantification of the phototaxis index (Figure S1B).

**Figure S1.**
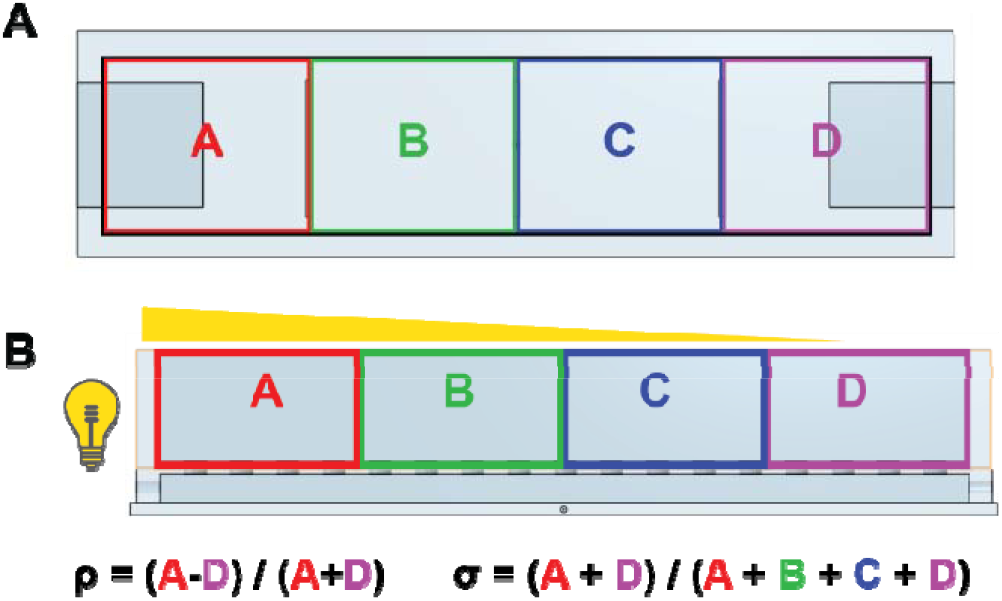
Phototaxis assay. A) Schematic of phototaxis chamber viewed from above. Small rectangles are demarcations for dividing the chamber into quarters. B) Schematic of phototaxis chamber viewed from the side with light coming from the left side. Methods for calculating phototaxis index (ρ) and phototaxis strength (σ).

For each experiment, between 100-200 cells were placed into the specimen area with enough water to fill the chamber up to an approximate depth of 4 mm. The chamber, with *Stentor* cells, was then placed on the stage of a dissection stereomicroscope (AmScope SM2-TZ) with a Thorlabs monochrome CMOS (CS135MUN; Thorlabs) in darkness and left in to acclimate for 1-2 minutes. For general phototaxis, a natural white LED, which covers a wide spectral band of light, was placed on one end of the chamber. For action spectrum analysis, LEDs of specific spectral wavelengths were used; these are [380 nm, 410 nm, 440 nm, 460 nm, 490 nm, 520 nm, 560 nm, 590 nm, 600 nm, 605nm, 625 nm, 660 nm, 730 nm, and 850 nm]. After dark acclimation, the LED was lit just bright enough so as not to light up the opposite ‘dark’ end of the chamber. Brightness was modulated by altering the voltage fed to the LED such that its Illumination (as measured using a lux meter held approximately 0.5 cm from the light source) ranged between 9,000 to 10,000 lux for all visible light LEDs and 500 to 1,000 lux for near infrared (nIR) and infrared (IR) LEDs. *S. stipatus* spec. nov. *cells* were then disturbed by pipetting up and down (to prevent attachment) and left for 5 minutes in the same dark environment with the LED on. After the 5-minute LED exposure, the LED was turned off and removed, the transillumination light on the dissection microscope was turned on and a quick video scan was taken of entire chamber using ThorCam software (Thorlabs). Videos were analyzed in FIJI (ImageJ); cells in each quadrant were counted and totaled. All of these experiments were performed between 12PM and 4PM to ensure a baseline strong positive phototaxis response.

Phototaxis was measured as a ‘phototaxis index’ by comparing the number of cells within the first (A, light) and fourth (D, dark) quadrants only (Figure S1B). Cells in the middle two quadrants were assumed to be either indifferent or weakly phototactic. This index was calculated as ⍰= (A - D) / (A + D). The strength of phototaxis was measured by comparing the total number of cells in A and D versus the total number of cells, σ = (A + D) / (A + B + C + D). The action spectrum was thus constructed by comparing the phototaxis index of *S. stipatus* spec. nov. cells exposed to the range of light wavelengths mentioned above.

### Diurnal expression of phototaxis

For the analysis of diurnal expression of phototaxis we performed phototaxis experiments as described before, but using only white light (∼5000K). A *S. stipatus* spec. nov. culture, grown under 12L/12D conditions, was assayed every hour from 4AM through 9PM. Following experimental light exposure, cells were put back into proper light or dark conditions (depending on the time of day) in a separate culture, ensuring no cells were re-used following disruption of their light-dark cycle.

### Directionality Analysis

Directionality was measured using the Directionality plugin in FIJI (ImageJ). Approximately 200 *S. stipatus* spec. nov. cells were placed in an imaging chamber and allowed to acclimate for 5 minutes. Following acclimation, the chamber was placed atop a low magnification dissection stereomicroscope (AmScope SM2-TZ) with a Thorlabs monochrome CMOS (CS135MUN; Thorlabs) and cells were agitated by pipetting up and down to prevent attachment. For uniform light conditions, the under-stage LED light of the microscope was turned on and the central part of the chamber was imaged for 1 minute at 5 frames per second using ThorCam software (Thorlabs). For asymmetric light conditions, the under-stage LED light of the microscope was turned as low as possible to enable normal imaging with the camera at 5 frames per second; a second brighter light source - a 3-watt natural white light (∼5000K; Chanzon) LED - was placed on the left side of the phototaxis chamber. The central part of the chamber was imaged for 1 minute at 5 frames per second. After imaging, 120 frames of each condition were used to generate Movie 2 and Movie 3 (played back at 12 frames per second). The same 120 frames were used to generate time projections. Due to the shape of our phototaxis chamber, as a long narrow rectangle, we chose to measure directionality in a central region of the resultant time projections away from the biasing effect of the walls which could redirect motile cells. We compared directionality under uniform lights versus asymmetric light using local gradients orientation with 180 angle bins (1 degree each).

### Centrifugation of *S. stipatus* spec. nov

Approximately 100 *S. stipatus* spec. nov. cells were pooled into a 1.5ml centrifuge tube in approximately 700 microliters of SPW. *S. stipatus* spec. nov. were centrifuged in a tabletop microcentrifuge at 12,500 RPM for 5 minutes. Following centrifugation, cells were allowed to recover for 1 minute before being transferred to an imaging chamber for microscopy. Imaging was performed using a low magnification dissection stereomicroscope (AmScope SM2-TZ) with a Thorlabs monochrome CMOS (CS135MUN; Thorlabs) and ThorCam software (Thorlab) at 10 frames per second.

## RESULTS

### Isolation and ecological description of *S. stipatus* spec. nov

*Stentor stipatus* spec. nov. was first identified from samples collected from a white cedar swamp which connects Sols and Jones ponds in the Two Ponds Conservation Area of Falmouth, Massachusetts in Cape Cod (approximate coordinates of 41°33’52.0”N 70°36’26.9”W) in late July. This *Stentor* species was found readily within the silt and decaying plant matter, and on bladderwort plants in the shallows of the swamp that connects Sols and Jones ponds. Both Sols and Jones ponds exhibit large blooms of *S. pyriformis* in the late spring through late summer (April through July); yet strikingly, despite connecting these two ponds, the white cedar swamp is generally devoid of *S. pyriformis*. The white cedar swamps of Cape Cod are known for being iron-rich (Motzkin, 1991), and indeed this swamp had significantly higher levels of iron than most other freshwater sources, at roughly 20-fold more iron than the second highest detected level (Table 1). *S. stipatus* spec. nov. primarily pooled in the sunlit regions of the swamp; due to iron and other particulates, the water was a dark orange-brown when collected, occluding initial spotting of free-swimming *Stentor* cells. However, after sediments settled and immediately upon inspection via low magnification stereomicroscope, hundreds of fast-swimming dark trumpet-shaped cells were readily seen pooling above the bright under-stage light of the microscope. Initial morphological characterization by low-magnification stereomicroscope confirmed these cells as an indeterminate *Ciliate*, likely a species of *Stentor*.

### Morphological description of *S. stipatus* spec. nov

By low magnification transmitted light microscopy, *S. stipatus* spec. nov. cells presented as dark brown to black and pyramidal, shaped almost like watermelon seeds (Figure 1A; Figure S2). Using higher magnification along with either oblique light or dark field microscopy, a two-toned coloration became more apparent. *S. stipatus* spec. nov. cells were predominantly green in color (Figure 1B-F; Figure S3A; Figure S4A) along their cortical cytoplasm with some patchy reddish-brown cytosolic pigmentation which appeared to cluster around their presumptive macronucleus (Figure 1B-F, Figure S4A, C). Neither transmitted light nor dark field imaging alone were sufficient to concretely identify either the position or shape of the macronucleus, but fluorescence imaging with DAPI nuclear signal (Figure S4D) verified the central location of a single spherical macronucleus deep within the cytoplasm in a pattern of localization consistent with the dark pigmentation. At higher magnification, cortical rows of alternating green and reddish-brown became visible (Figure 1F). Upon further investigation, it became apparent that the *Stentor* cells were full of cortically-proximal rows of microalgal cells (Figure S4A) which were the source of the cortically enriched green color. Further, these microalgal cells were extremely autofluorescent (Figure S4B) allowing us to fix and image stentor cells and their endosymbionts with a high-powered fluorescence microscope. Closer inspection of the distribution of the algal cells revealed that they are densely packed within the cortex, with few to no algal cells present in the internal cytoplasm. Cortically packed algae are uniformly distributed everywhere along the cortex except the region around the membranelles of the cell (Movie 1, visible gap in Figure S4B).

**Figure 1.**
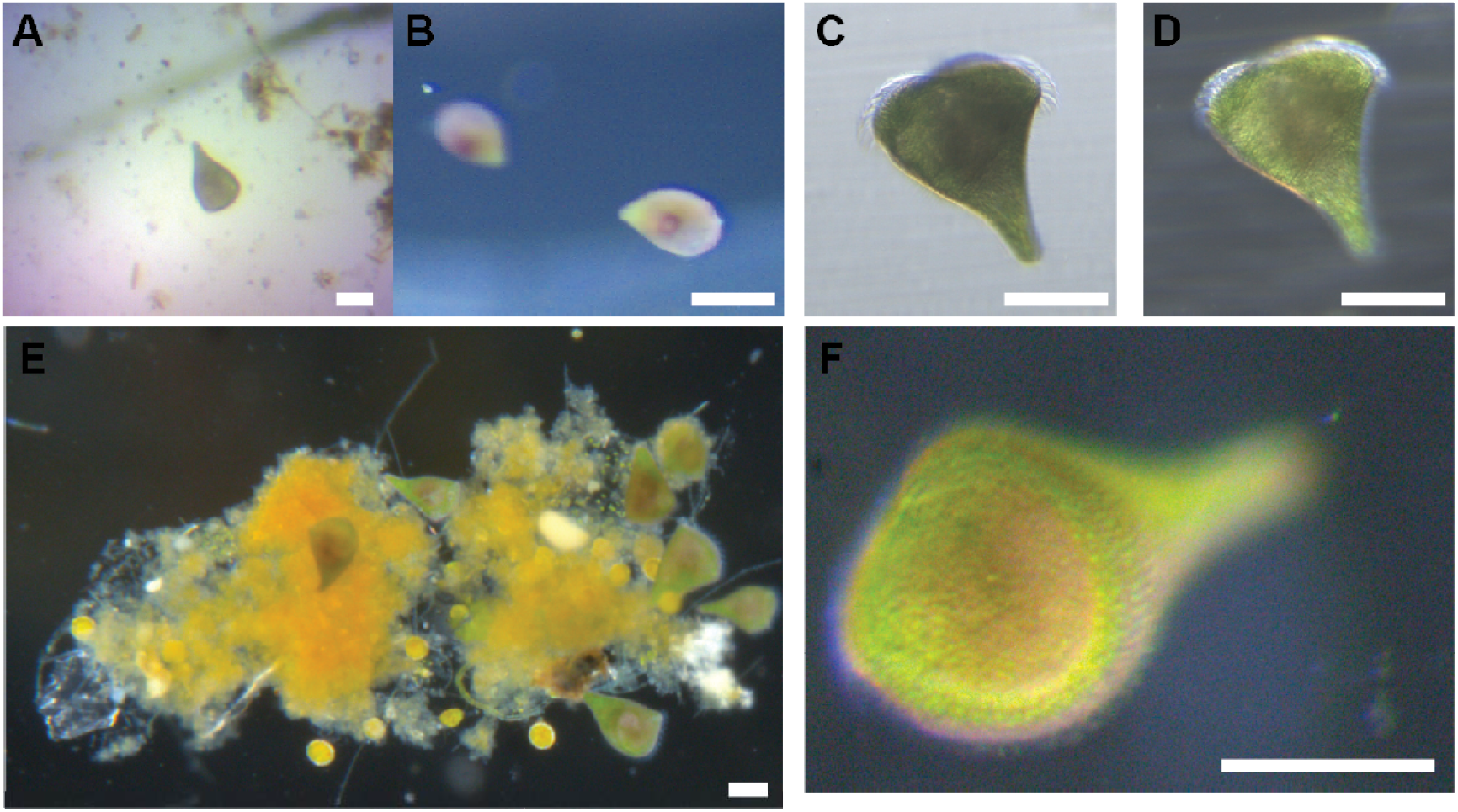
*Stentor stipatus* spec. nov. are densely packed with algae and pigment aggregates. A) By low magnification transmitted light imaging, *S. stipatus* spec. nov. cells appear dark brown to black and seem to have dark pigment granule aggregate inclusions. B) By low magnification oblique light imaging, *S. stipatus* spec. nov. cells demonstrate clear green and reddish-brown pigmentation with dense patches of brown color located in the internal cytoplasm. C) Higher magnification transmitted light image showing green color with dark brown aggregates. D) Same cell as in C, imaged via darkfield, dark brown aggregates are now apparent as reddish-brown pigmentation. E) Low magnification darkfield image of a cluster of *S. stipatus* spec. nov. cells adhering to organic substrate. All cells have visible reddish-brown cytosolic pigmentation. F) Higher magnification darkfield image of a *S. stipatus* spec. nov. cell. Cortical striations of alternating green and reddish-brown visible laterally along the cell. Scale bars are approximately 100 microns.

**Figure S2.**
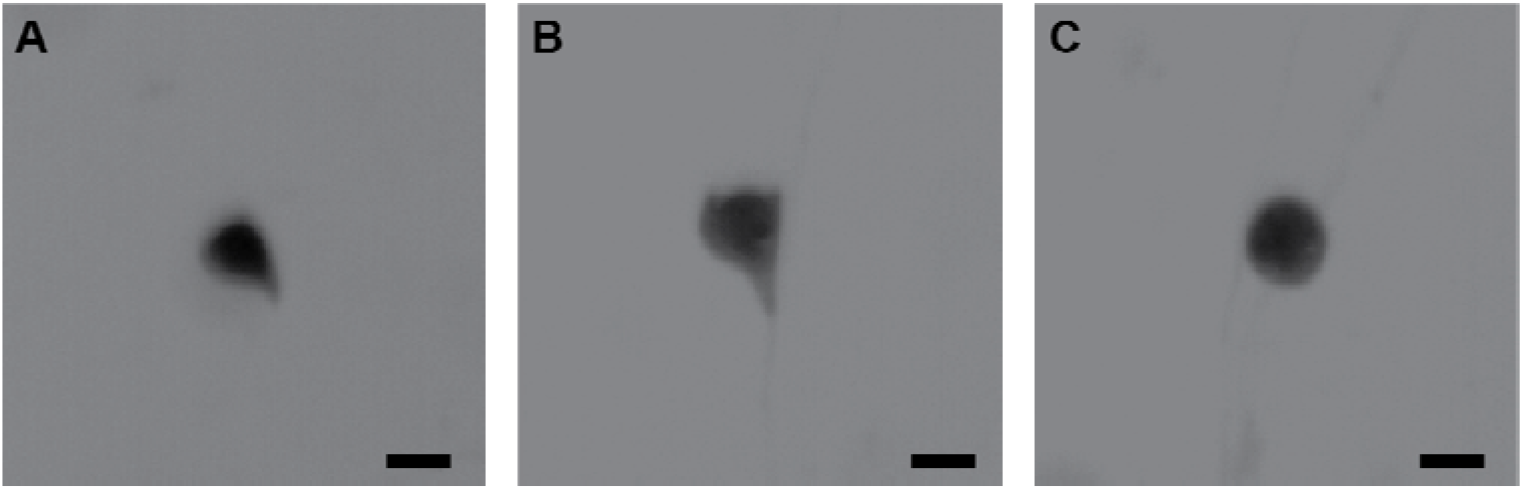
*S. stipatus* spec. nov. cell shape and contraction. A) Image of a swimming S. stipatu spec. nov. cell, demonstrating the hydrodynamic ‘tear drop’ shape. B) Image of a cell attached to a substrate demonstrating the pyramidal stout shape. C) Image of the same cell in B after mechanical stimulation-induced contraction. Scale bars are approximately 100 microns.

**Figure S3.**
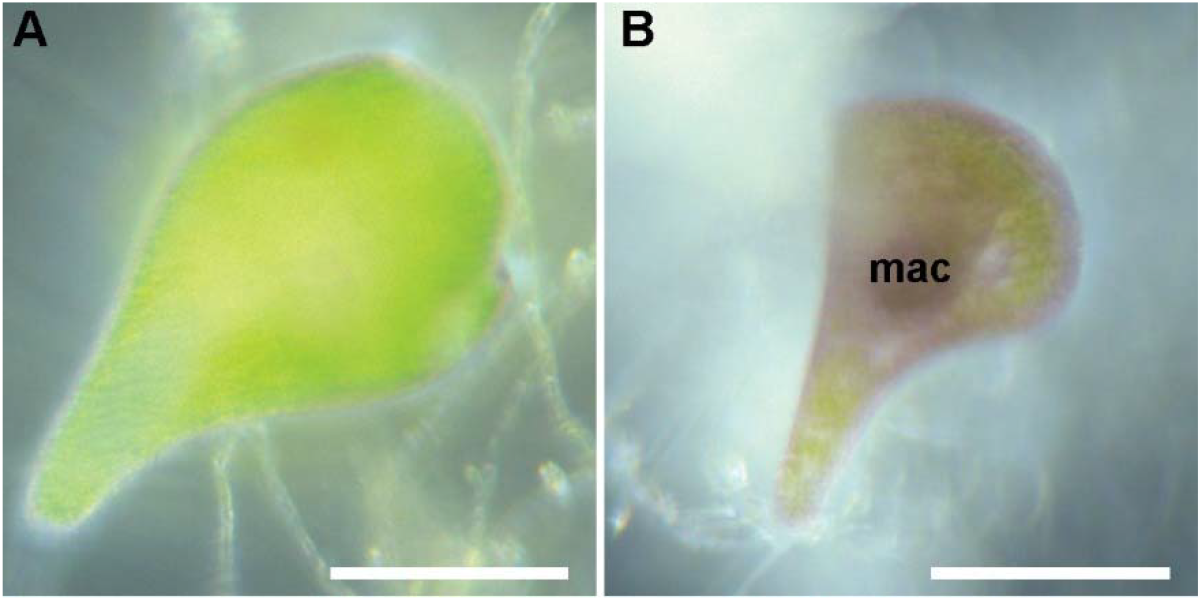
Coloration of *S. stipatus* spec. nov. compared to *S. amethystinus*. A) High magnification darkfield micrograph of an *S. stipatus* spec. nov. cell with predominantly green cortical coloration, slight reddish-brown pigmentation is visible and is present in the cytosol. B) High magnification darkfield micrograph of presumptive *S. amethystinus* cell showing predominantly reddish-brown pigmentation throughout the cell with a darker density around the macronucleus (mac). Both images were taken under the same imaging conditions. Scale bars are approximately 100 microns.

**Figure S4.**
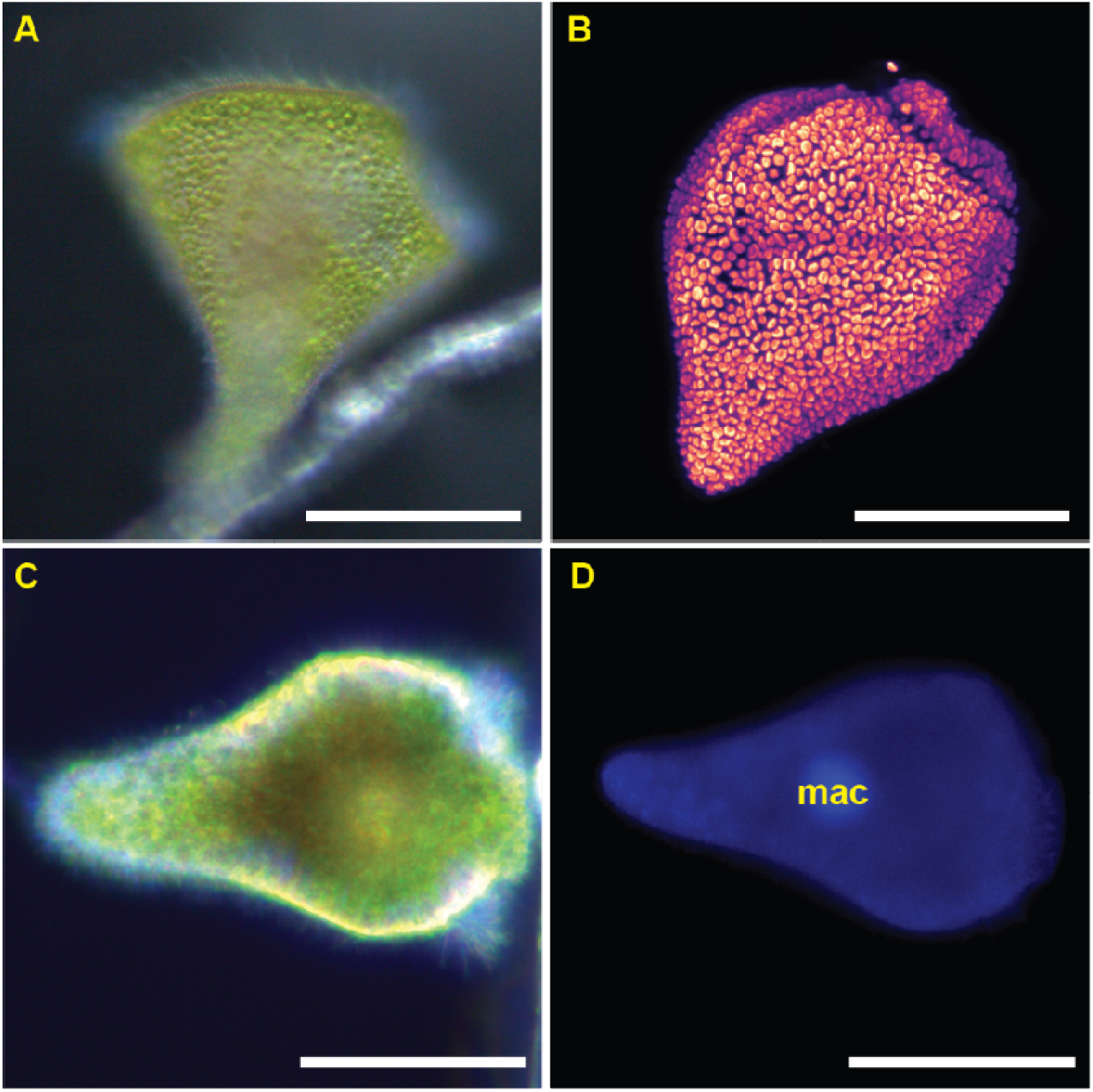
*S. stipatus* spec. nov. has a cortical enrichment of endosymbiotic algae. A) High magnification darkfield micrograph of a *S. stipatus* spec. nov. cell highlighting the dense cortical packing of microalgae within the cell. B) High magnification fluorescence micrograph of a *S. stipatus* spec. nov. cell showing autofluorescence of endosymbiotic algae. C) High magnification darkfield micrograph of *S. stipatus* spec. nov. cell highlighting the dense reddish-brown pigmentation which appears to surround the macronucleus. D) Autofluorescence and DAPI fluorescence signal of the same cell in C with the macronucleus labeled (mac). Scale bars are approximately 100 microns.

Initial morphological assessment suggested our isolate from the white cedar swamp could be a variant of *S. amethystinus*, though *S. amethystinus* has largely been described as having a vivid and more uniform red pigmentation (Foissner and Wölfl, 1994; Höfle *et al*., 2014) than what we observed for *S. stipatus* spec. nov. cells. Indeed, another strain, more recently isolated from a pond in Flat Rock, North Carolina, which has anecdotally been reported to more closely match *S. amethystinus* morphology, demonstrated a clear difference in coloration when compared to *S. stipatus* spec. nov. cells (Figure S2A, B). As seen by high magnification dark field microscopy, *S. stipatus* spec. nov. cells generally carry much less red pigmentation than cells of the presumptive *S. amethystinus* variants (Figure S1A, B) which appear to have bold reddish-brown pigmentation throughout along with a highly pigmented concentration around their presumptive macronucleus.

*S. stipatus* spec. nov. cells were found to be very motile; these cells often attach to plant substrate by their holdfast but readily detach and freely swim around following minimal perturbation. The motile shape of these cells is teardrop-like, with cells approximately 197±11.8 micrometers in length by 134±11.3 micrometers in width, with an aspect ratio of approximately 1.48±0.13 (n = 32 cells; Figure S2A). At full extension, these cells did not change in shape much, with a slight decrease in aspect ratio to 1.4±0.2 (n = 35 cells; Figure S2B) as cells appeared to expand slightly more in width than length. Fully extended, sessile cells had a large, flat frontal field, with membranelles exhibiting a stereotype metachronal beat pattern which was observable by transmitted light with low magnification. Overall, this *Stentor* species exhibits limited expansion between swimming and sessile forms with statistically similar aspect ratios (Figure S2A, B), making it similar to *S. pyriformis* in this regard (Hoshina *et al*., 2021). Upon contraction, cells withdrew their membranelles and shrank to an approximate aspect ratio of 1.0±0.1 (n = 11 cells), nearly spherical in shape (Figure S3C).

### S. *stipatus* spec. nov. is phylogenetically distinct from *S. amethystinus*

By morphological description alone, *S. stipatus* spec. nov. could be a variant strain of *S. amethystinus*. We therefore set out to describe the molecular phylogenetic relationship between *S. stipatus* spec. nov. and other known *Stentor* species. Phylogenetic work based on analysis of 18S SSU rDNA sequence has clarified our understanding of the genus and validated a few suspect species and species complexes (Thamm *et al*., 2010). We therefore used 18S SSU rDNA analysis to test whether *S. stipatus* spec. nov. was indeed a distinct species from its morphologically-similar relative *S. stipatus*. The maximum likelihood (ML) tree generated with our *Stentor* sequences was found to be in general agreement with previously generated trees of the genus (Thamm *et al*., 2010; Hoshina *et al*., 2021), preserving the general relationships between established *Stentor* species. Within our ML tree, *S. stipatus* spec. nov. sits branching off the same family as *S. pyriformis* and *S. amethystinus* and is separated from *S. amethystinus* by 0.0177 nucleotide substitutions per site, comparable to the separation between *S. amethystinus* and *S. pyriformis* (0.0175), or *S. tartari* and *S. elegans* (0.0127) (Figure 2). Furthermore, bootstrapping (90.7), single-branch Bayesian inference (0.987), and SH-aLRT (89) all strongly support the monophyly of the *S. stipatus* spec. nov. branch (Figure 2).

**Figure 2.**
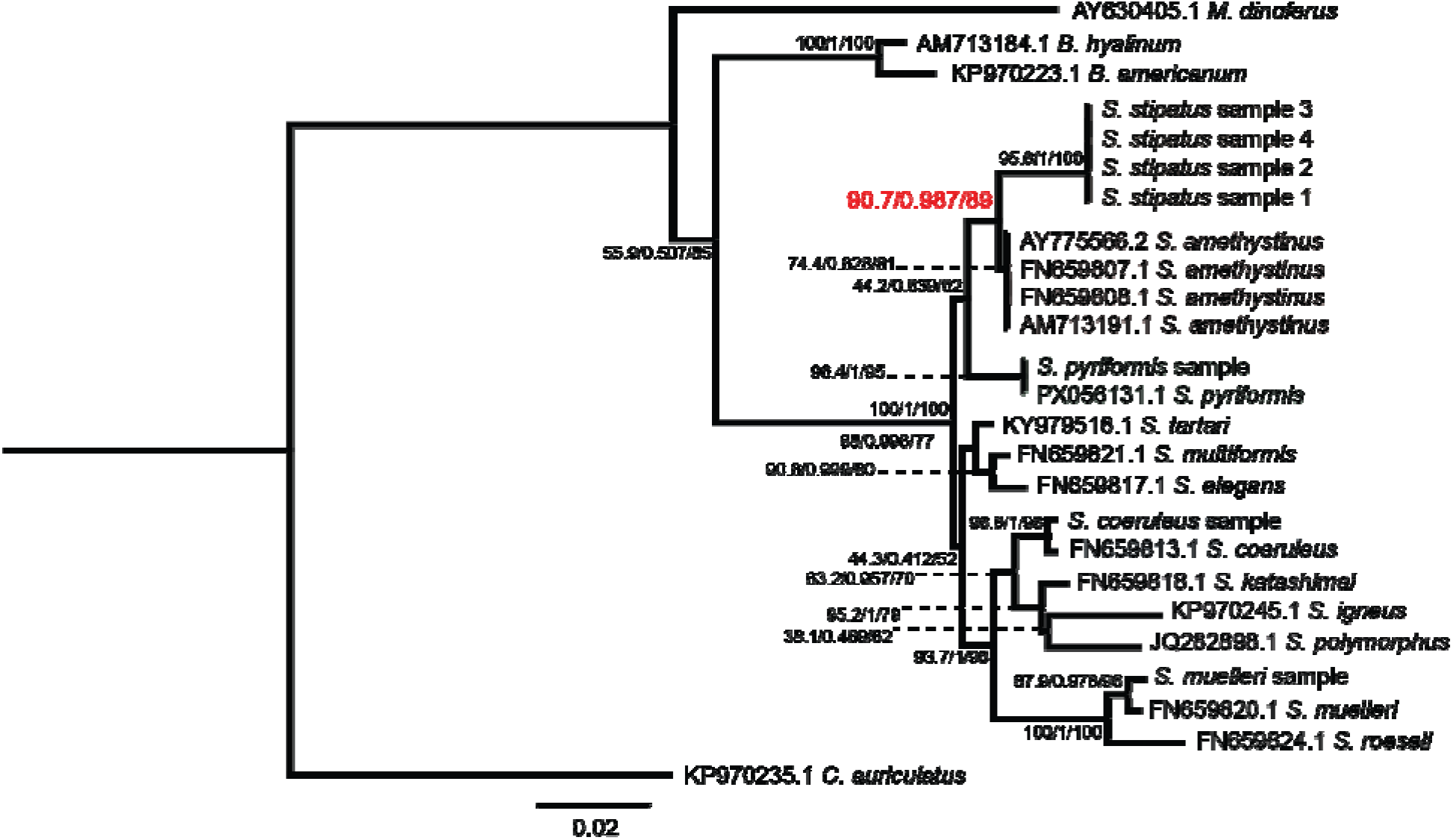
*S. stipatus* spec. nov. is phylogenetically distinct from other *Stentor* species. A) Maximum likelihood (ML) tree estimate, with branch lengths showing nucleotide substitutions per site. Nodes show reliability measure scores (UfBoot/ aBayes / SH-aLRT ). Red text highlights the node between *S. stipatus* spec. nov. and *S. amethystinus*. Major families are highlighted by colored boxes, *S. stipatus* spec. nov. is highlighted in the brown box. Scale bar indicates nucleotide substitutions per site.

### *S. stipatus* spec. nov. cells habituate to mechanical force

*S. stipatus* spec. nov., like all other members of the genus, discernably contracts when presented with unfavorable external stimuli (Tartar, 1961) including mechanical stimulation by jarring or poking. Contraction is a specifically elicited behavioral response which *Stentor* do not randomly enact under standard culturing or imaging conditions, and habituation to repeated mechanical stimulation-induced contraction has been described carefully for *S. coeruleus* before (Wood, 1988; Rajan *et al*., 2023b). We therefore set out to characterize the contraction response of *Stentor stipatus* spec. nov. cells after repeated mechanical stimulation and whether habituation. Indeed, we found that *S. stipatus* spec. nov. cells habituate within an hour to high-force stimuli presented as described before (Figure 3; (Rajan *et al*., 2023b)) as evidenced by the drop in population contraction following stimulation, which is not seen in cells not exposed to repeated stimulation. *S. coeruleus*, on the other hand, typically do not habituate to such high-force stimulation within the same time frame (Figure 3). *S. stipatus* spec. nov. cells also appeared to be less mechanically sensitive than *S. coeruleus* cells given that the prescribed high-force stimulus as presented caused 100% of *S. coeruleus* cells to contract but only elicited approximately 75% of *S. stipatus* spec. nov. cells to contract.

**Figure 3.**
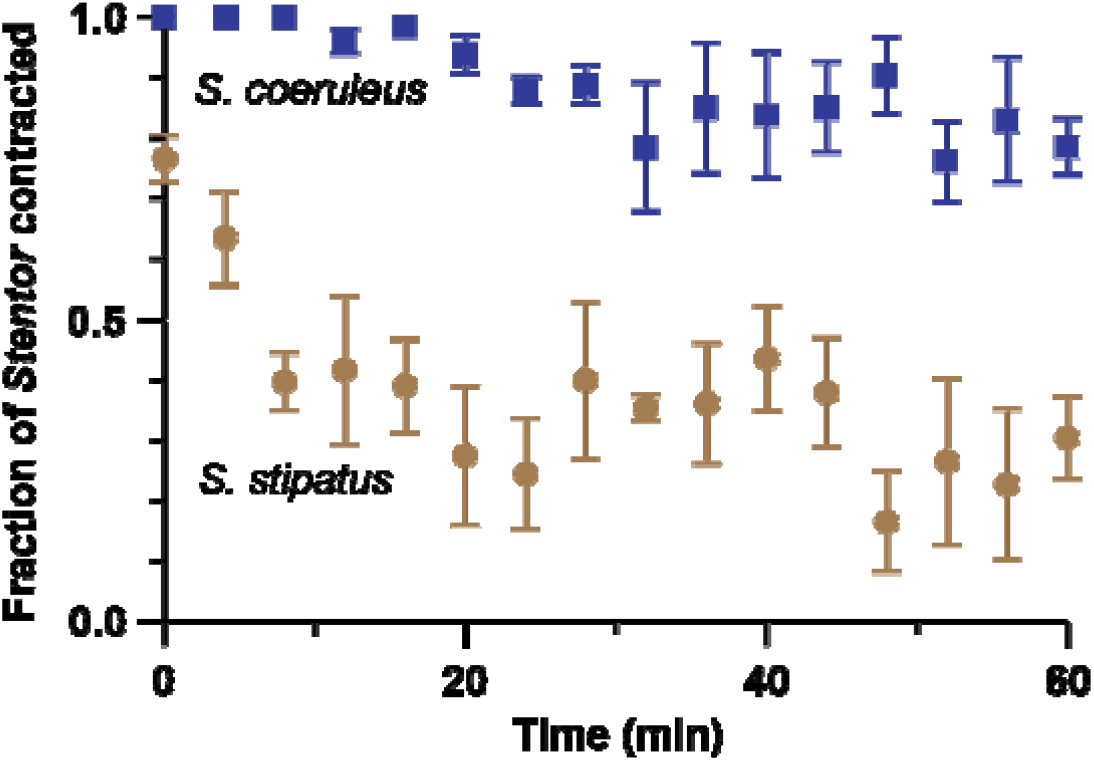
*S. stipatus* spec. nov. habituates to high force taps. (Brown circles) Habituation in *S. stipatus* spec. nov. cells (3 replicates per data point; n = 6-32 cells per replicate). (Blue squares) habituation in *S. coeruleus* cells (3 replicates per data point; n = 6-67 cells per replicate). Level 3 taps; 1 tap / minute. Error bars represent standard error of the mean. n values represent the range for the number of cells that were anchored and within the microscope field of view for each series of replicates.

### Action Spectrum analysis of *S. stipatus* spec. nov. reveals multiple phototaxis peaks

Initial isolation of *S. stipatus* spec. nov. cells from water samples revealed clear positive phototaxis, thus we set out to characterize this light response. On the short timescale, *S. stipatus* spec. nov. cells exhibited strong positive phototaxis when presented with asymmetric broadband white light (∼5000K) or red-shifted light (380-840 nm) light, with a phototaxis index of 0.75±0.23 (n = 3 experiments with a total of 413 cells) and 0.70±0.24 (n = 3 experiments with a total of 476 cells) respectively. The overall strength of this phototactic response (the percentage of total cells in either of the two extreme quadrants in our chamber) was measured at 75% for white light and 76% for red-shifted light.

We next explored the action spectrum of *S. stipatus* spec. nov. and found positive phototaxis for all measured wavelengths with the phototaxis index varying between wavelengths. The phototactic response with highest phototactic index appeared to be around 590-600 nm and the weakest around 850 nm; a secondary, smaller peak in phototactic activity appeared under exposure to 520 nm light. A slight dip in activity around 490 nm also suggests a potential third, much less defined peak in the blue-green range of light around 430 nm (Figure 4A). It is worth noting that for all positive phototaxis recorded, motion towards light was extremely directed towards the light source (Movie 3; Figure 4D) rather than random movement (Movie 2; Figure 4C) agnostic to the light source leading to eventual accumulation.

**Figure 4.**
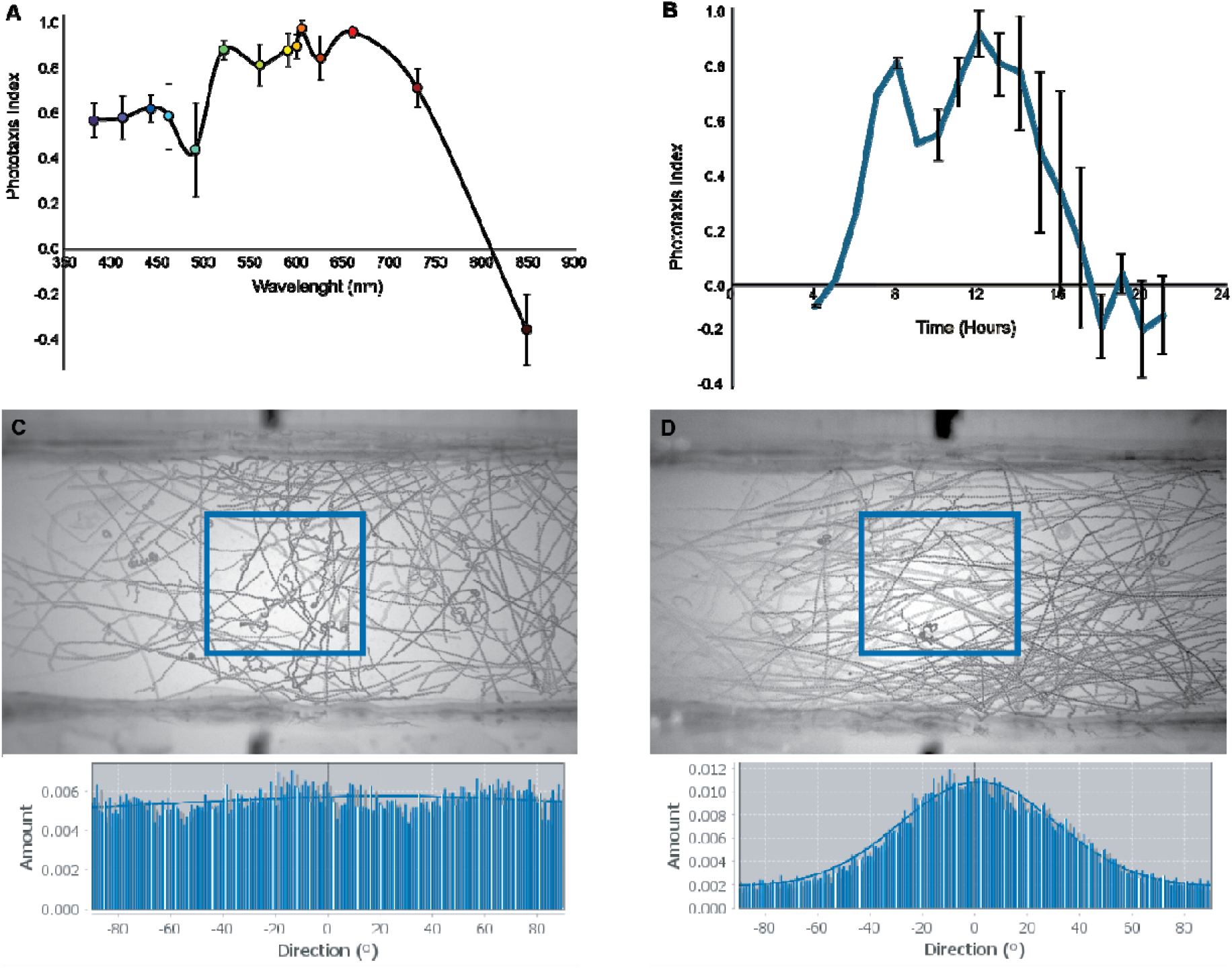
Action spectrum and phototaxis response of *S. stipatus* spec. nov. A) Graph shows the phototaxis response of *S. stipatus* spec. nov. to different wavelengths of light ranging from ultraviolet (380 nm) to infrared (850 nm). For each experiment, between 120-250 cells were placed in the phototaxis chamber and counted; the experiment was repeated 4-5 times for each wavelength measured. B) Graph shows the phototaxis response of *S. stipatus* spec. nov. to ∼5000K white light at different times of the day with time of light-dark transition (yellow versus blue areas in graph). For each experiment, approximately 200 cells were placed in a phototaxis chamber; the experiment was repeated 4 times for each time point. Errors bars demonstrate standard error of the mean. C) Top: Time projection of *S. stipatus* spec. nov. cells swimming in uniform light, blue inset shows region used for directionality analysis. Bottom: Graph shows histogram of directionality detected, demonstrating random swimming. D) Top: Time projection of *S. stipatus* spec. nov. cells swimming in asymmetric light focused on the left side of the chamber, blue inset shows region used for directionality analysis. Bottom: Graph shows the histogram of directionality detected, demonstrating directed swimming towards the light source.

### *S. stipatus* spec. nov. demonstrates a cyclical pattern in phototactic response strength

There is evidence that some species of *Stentor* may have variable phototaxis response throughout the day (Boudreau *et al*., 2025). In our initial efforts to collect *S. stipatus* spec. nov. cells from their source we noted that (1) cells were much more likely to be motile if isolation were carried out during daytime hours, and (2) cells could easily be isolated by their positive phototactic response - but only during daylight hours. In the evening, *S. stipatus* spec. nov. cells appeared to transition into a less motile behavioral pattern, preferring to anchor within substrate or debris, and no longer pooling as strongly around strong light. We therefore wanted to know whether the general phototactic response of this species varied in a diurnal manner. Indeed, the phototactic response of cells cultured under 12L/12D conditions (7AM and 7PM transition of lighting) around 4AM is negligible (-0.1) and rapidly increases to a peak by 12PM (0.91), before diminishing again by 6PM (-0.17) and remaining negative thereafter (Figure 4B).

### Centrifugation of *S. stipatus* spec. nov. affects phototaxis

*S. pyriformis*, which contains hundreds of endosymbiotic microalgae of genus Chlorella, expels its microalgae following high-speed centrifugation, after which *S. pyriformis* phototaxis is significantly reduced (Boudreau *et al*., 2025). Since *S. stipatus* spec. nov. demonstrates both dark pigmented patches along with peripheral green microalgae, we decided to test whether pigmentation or endosymbionts could be removed by centrifugation, and whether this would influence the phototaxis demonstrated by *S. stipatus* cells. Following centrifugation, *S. stipatus* spec. nov. did not appear to lose neither pigmentation nor algae but rather demonstrated significant redistribution of total pigmentation by low magnification transmitted light imaging, such that half of the *Stentor* (divided along the long axis; Figure 5A) contained both green and red pigmentation while the other half was entirely transparent. In low light conditions, or with uniform light, these *Stentor* cells demonstrated no other significant changes in morphology, behavior, or motility following centrifugation. If bright asymmetric light was applied to centrifuged *S. stipatus* spec. nov. cells, however, they exhibited altered swimming behavior; rather than swimming directly toward bright light as shown before, centrifuged cells swam in circles generating little effective movement (Movie 5; Figure 5A’). Over the course of about 10 minutes, the pigmentation in centrifuged cells was seen to redistribute such that they were eventually fully covered by dark coloration as before centrifugation (Movie 6; Figure 5B) and swimming became normal again (Movie 7). The demonstrated change in swimming behavior could have been partly due to a redistribution of mass if the dark pigmentation were more dense than non-pigmented cytoplasm. However, given that the dark pigmentation of *S. stipatus* spec. nov. cells is the result of a combination of reddish pigmented patches and green microalgae, we decided to fix and image cells immediately after centrifugation to see whether algae were indeed redistributed in a way that could explain the change in phototactic response. Fixed centrifuged cells demonstrated a complete redistribution of the internal microalgae such that all algal cells were still cortical but were entirely within the dark pigmented ‘half’ of the cortical cytoplasm (Figure S2C; Movie 4), providing some potential explanation for why centrifuged *S. stipatus* spec. nov. cells change their swimming behavior in asymmetric light - namely, that the algae was involved in either detecting the light, or modulating swimming behavior, or both.

**Figure 5.**
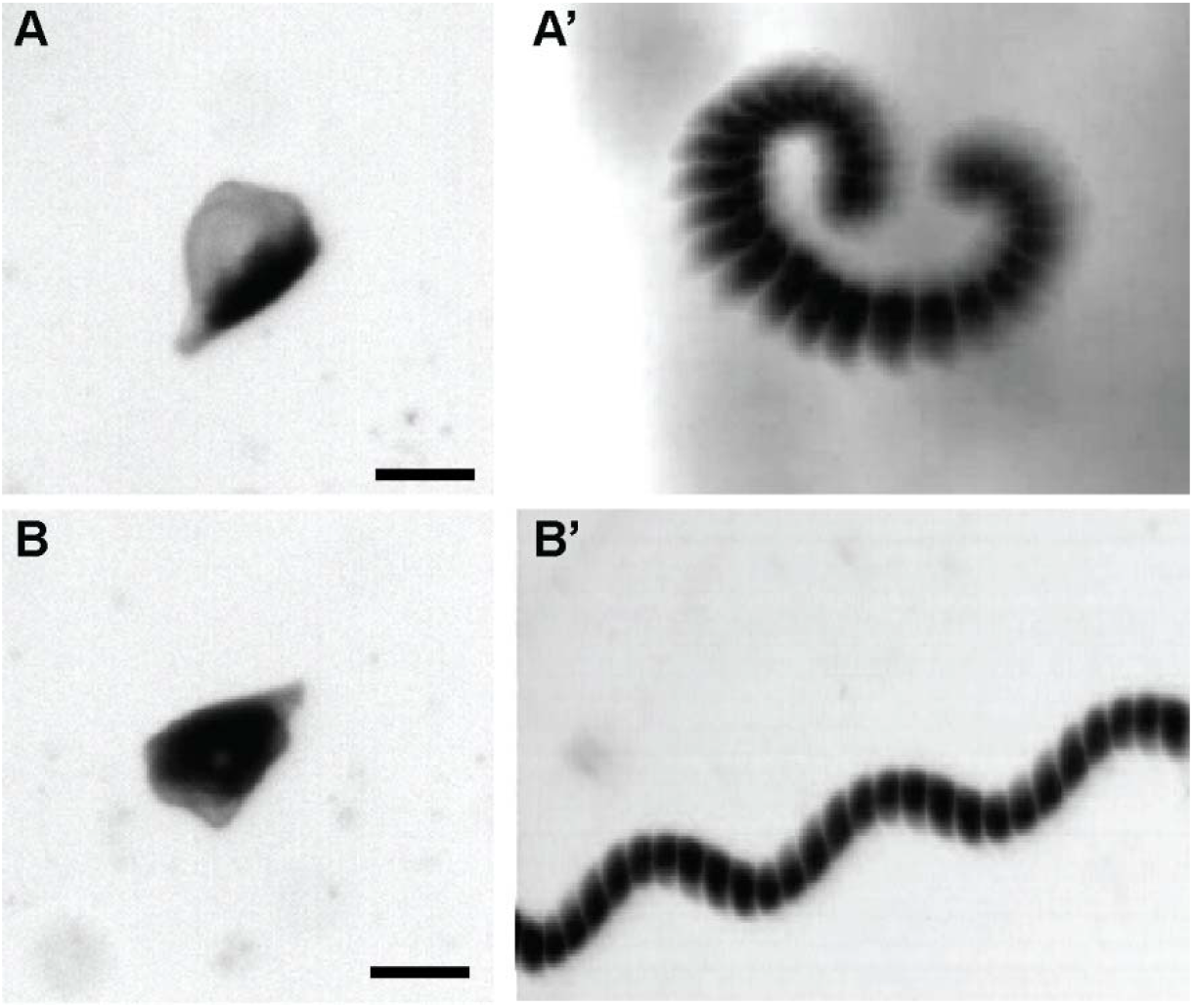
Effects of centrifugation on *S. stipatus* spec. nov. motility and light response. A) Close-up of *S. stipatus* spec. nov. cell following high-speed centrifugation. Dark pigmentation redistributes in consistent left-right light-dark pattern shown in all assayed cells (n = 22). A’) Time projection of swimming centrifuged *S. stipatus* spec. nov. cell. When presented with asymmetric light, these cells swim in a less processive manner than normal (3/3). B) After several minutes of recovery, dark pigmentation is redistributed throughout the cell (n = 10). B’) Time projection of swimming centrifuged *S. stipatus* spec. nov. cell after recovery. Following recovery of normal pigment distribution, cells now swim in the normal corkscrew pattern. Image squashed horizontally to fit figure (3/3). Scale bars are 100 microns.

## DISCUSSION

Here, we introduce *Stentor stipatus* spec. nov. as a new species based on molecular phylogenetic analysis (Figure 2), which reveals *S. stipatus* spec. nov. as a monophyletic clade separate from its nearest relative, *S. amethystinus*. UfBoot, along with aBayes and SH-aLRT analysis, all support a tree topology where *S. stipatus* spec. nov. samples form a single cluster that branches apart from *S. amethystinus* (Figure 2). Interestingly, these three methods would suggest that *S. stipatus* is not entirely monophyletic or could be subject to significant variability (based on values between PX056133 and the rest of the group) (Figure 2). However, when considered along with nucleotide substitutions per site (Figure 2), which also supports separation from *S. amethystinus* but does not distinguish between members of the *S. stipatus* spec. nov. group, these results are consistent with *S. stipatus* spec. nov. being a monophyletic species.

While 18S SSU rDNA sequence comparison is a widely used method for construction of phylogenies, single gene phylogenies can often fail to capture the species phylogenetic relationships due to incomplete lineage sorting or introgression, requiring multiple gene sequence comparison (Chen *et al*., 2015; Fernandes *et al*., 2016). Thus, even though we present strong evidence which meets the genus standard for the definition of *S. stipatus* spec. nov. as a discrete species, we cannot rule out that species tree topology may be discordant with the 18S SSU gene tree. To better differentiate between *S. stipatus* spec. nov. and *S. amethystinus*, we will need to obtain additional genetic material from both purported species for more detailed sequence comparisons. One limitation of taxonomical research on protists is inadequate environmental sampling and re-isolation (Adl *et al*., 2007), which can lead to a paucity of high-quality genetic material for phylogenetic comparisons, resulting in clades with low bootstrap values across the *Stentor* genus (Figure 2). Within the *Stentor* genus, the *S. amethystinus* clade is particularly disorganized due to variety in available sequences and is sometimes not even considered monophyletic (Fernandes *et al*., 2016). Once additional isolates are gathered from field samples, molecular barcodes using specific genes can be used to distinguish between species more finely, as has been done in other protist genera such as *Tetrahymena* (Doerder, 2019).

Indeed, morphological comparison denotes how closely related *S. stipatus* spec. nov. is to *S. amethystinus. S. amethystinus* is not reported to have the same coloration that we have observed in *S. stipatus* spec. nov., but both are in a comparable size range (Heep *et al*., 1998). The dark coloration of both *S. stipatus* spec. nov. and *S. amethystinus* appears to be due to a combination of the green microalgae within the cell as well as some form of reddish pigmentation. While our phototaxis action spectrum data suggest that *S. stipatus* spec. nov. may also contain a photopigment like other pigmented *Stentor*, it is not entirely clear whether the reddish-brown pigmentation of *S. stipatus* spec. nov. is contained in the form of pigment granules like in *S. amethystinus*. Furthermore, both *S. stipatus* spec. nov. and *S. amethystinus* are less contractile (Figure 3; (Cragg, 1971)) than other members of the *Stentor* genus– though this could be an artifact of their similar rounded morphology. Nevertheless, morphology alone is not an adequate identifier of *Stentor* species because *Stentor* morphology varies according to the region where conspecific cells were isolated (Heep *et al*., 1998).

Our initial characterization shows that *S. stipatus* spec. nov. cells contain a densely packed shell of algae which provide the predominant green color of this strain (Figure S4). We have not yet succeeded in isolating or culturing the algae to identify it and compare it to the known endosymbiotic algae of *S. pyriformis* (Boudreau *et al*., 2025). Furthermore, it is as yet unclear whether the secondary, reddish-brown coloration of *S. stipatus* spec. nov. is a form of amethystin or a similar pigment concentrated within cortical (and macronucleus-adjacent) pigment granules or something else entirely. One intriguing possibility is that pigmentation in *Stentor* could serve as a form of ‘sunblock’ meant to reduce the occurrence of UV light-induced DNA damage. Some *Stentor* species, which have gained endosymbiotic algae do not appear to have a need for pigmentation as their algal partners effectively shield their macronucleus from light, these species therefore have no visible pigment granules (*S. pyriformis* and *. polymorphus*). Others, like *S. amethystinus* and *S. stipatus* spec. nov. have retained pigmentation despite gaining algal endosymbionts. In the case of *S. stipatus* spec. nov. at least, it appears that the algae are tightly associated with the cortex of the cell while most of the pigmentation occurs internal to the algae - ensuring algal cells have access to light while also ensuring protection for the macronucleus. The differences between *S. stipatus* spec. nov. and *S. pyriformis*, which together bloom in the geographically adjacent though chemically distinct waters in Cape Cod at similar times of the year, could provide novel insights into unicellular physiology and adaptation. Similarly, studying the mechanisms by which *S. stipatus* spec. nov. regulates the localization of its pigments and algae could provide some new insights into complex cytoskeletal organization and dynamics. In the future we could centrifuge *S. stipatus* spec. nov. cells and then bisect centrifuged cells to help us isolate the reddish-brown patches of pigmentation within *S. stipatus* spec. nov. which could then be further analyzed by mass spectrometry. If these indeed contain a form of pigment it would be interesting to compare the biochemistry of this to the known pigments of other *Stentor* species including *S. coeruleus* and *S. amethystinus*.

The waters of the white cedar swamps in the Two Ponds Conservation Area in Falmouth are rich in iron to a level that appears lethal to closely related species of *Stentor* as we noted that *S. amethystinus* isolated from North Carolina could not survive for more than a few days in water mimicking the chemical composition of white cedar swamp water. Further, despite blooming in both Jones and Sols ponds, which are physically linked by the white cedar swamp, neither *S. pyriformis* nor *S. coeruleus* survive long in synthetic white cedar swamp water. Indeed, while some *S. pyriformis* have been collected from the same water samples as *Stentor stipatus* spec. nov., these *S. pyriformis* cells were generally unhealthy in appearance. It is therefore a possibility that the reddish-brown pigmentation in this *Stentor* species could be the result of incorporation or sequestration of heavy metal pollutants from their environment. At least one other major ciliate, *Tetrahymena*, has a demonstrated capacity to accumulate arsenic from its environment and even metabolize it into less toxic forms (Yin *et al*., 2011; Xiong *et al*., 2024). Furthermore, Chlorella has also been demonstrated to accumulate arsenic from its environment (Alharbi *et al*., 2023), making *S. stipatus* spec. nov. and other *Stentor* species, potentially capable of partial adaptation to heavy-metal contaminated waters. Further work to establish whether *S. stipatus* spec. nov. or other *Stentor* species are tolerant to heavy-metal contamination is ongoing.

Many characterizations of new species focus on morphological descriptions (Foissner and Wölfl, 1994), but our phototaxis and habituation experiments reveal that behavioral complexity is an essential feature of *S. stipatus* spec. nov. which can also distinguish it from other species. *S. stipatus* spec. nov. cells exhibit strong positive phototaxis over short timescales (Figure 4). Other *Stentor* species such as *Stentor pyriformis* exhibit positive phototaxis driven by the presence of obligate endosymbionts. S pyriformis cells cannot survive long without their algal partners so they logically maintain strong phototaxis in healthy cultures. *S. stipatus* spec. nov. may not require algae for survival (something else that needs to be tested if the algal partner can be removed), so differences in dietary needs might affect phototaxis in *S. stipatus* spec. nov. as well. Future experiments will thus require adoption of standard culturing methods, something which is not yet possible as we work towards identifying ideal culturing conditions. It is worth noting that recent attempts to culture *S. stipatus* spec. nov. in full dark conditions have proven unsuccessful, suggesting that perhaps their algal partner is necessary, though other reasons for culture failure cannot yet be ruled out.

The phototactic response of *Stentor* cells can be variable between different strains of the same species (Tartar, 1961) but it can also vary by time of day in some cases. Here, we show evidence for a cyclical pattern in the phototactic response of *Stentor stipatus* spec. nov. It should be noted that, while these data are not definitive, there is strong evidence within our results that phototaxis could be a behavior under circadian regulation. If the change in phototaxis response strength was merely diurnal - a simple response to external change in light conditions - we would expect to see the drop in phototaxis strength occurring sometime after the shift from light to dark conditions (7PM) or from dark to light (7AM). Rather, our data show a gradual loss in phototaxis strength throughout the day culminating in a total loss that generally occurs around one to two hours before the transition of lighting conditions in our cultures. This suggests that *S. stipatus* spec. nov. cells might regulate their phototactic response using an internal clock mechanism. Circadian rhythms have previously been demonstrated for phototaxis in other protists, including related ciliates like *Tetrahymena* (Wille and Ehret, 1968). Furthermore, several species of *Chlorella*, the genus of algae found within *S. pyriformis*, also demonstrate hallmarks of a circadian rhythm. Therefore, it should not be surprising that *S. stipatus* spec. nov., and perhaps other *Stentor* species, could have some form of circadian clock that regulates key behaviors. We present *S. stipatus* spec. nov. as an organism within which circadian clock machinery could be discovered and studied. First, and foremost, *S. stipatus* spec. nov. has both a potential algal endosymbiont as well as potential pigment granules, both mechanisms by which the cell can detect light. Our action spectrum analysis of *S. stipatus* spec. nov. demonstrates a *Stentor* which is adapted to detect light in both blue-green ranges like *S. pyriformis*;(Boudreau *et al*., 2025) and red–far red ranges like *S. coeruleus*; (Wood, 1976). Furthermore, the night-time shift to weakly negative phototaxis, rather than simply weakening of the positive phototaxis seen during the day, suggests not just a change in phototactic response strength but perhaps a directional shift in phototaxis behavior between day and night conditions. This could explain our observation that at night *S. stipatus* spec. nov. cells demonstrated a tendency to hide in substrate making isolation from collected samples harder.

One interesting feature of habituation in *Stentor*, similar to habituation in animals, is that cells habituate more rapidly to weaker stimuli. The fact that *S. stipatus* spec. nov. cells appear to be less sensitive (i.e. a smaller fraction of untrained cells contract in response to a given mechanical stimulus) and are also quicker to habituate (Figure 3) may provide an opportunity to investigate the relation between sensitivity and habituation rates. For example, one concrete question is whether the habituation curve of *S. stipatus* spec. nov. matches the habituation curve seen in *S. coeruleus* when different forces are used in the two species that give the same initial contraction probability. *S. stipatus* spec. nov. might also be a convenient organism for long-term time lapse imaging because it is less likely to contract in response to extraneous vibrations under the microscope. Genomic sequence comparisons among *Stentor* species might help to reveal the genetic basis for differences in mechanical sensitivity.

Further characterization of *S. stipatus* spec. nov. will include genome sequencing and phylogenomic comparison to other sequenced *Stentor* genomes. Currently, a major hurdle to this goal remains in that while we have succeeded in establishing stable culturing methods for this species, we have yet to grow these cells well enough to establish stable clonal cultures of high enough density for deep genome sequencing via next generation sequencing techniques. Once we further refine and identify the optimal conditions for long-term *S. stipatus* spec. nov. culturing in the lab, we can also carry out additional replicates for phototaxis and habituation experiments. In particular, we can test the stimulus-specificity and force-specificity of habituation in *S. stipatus* spec. nov. to check if they match the patterns seen in wild-caught *S. coeruleus* cells, which can habituate to other non-mechanical stimuli such as light (Wood, 1973). A current limitation of our habituation experiments is that while the *S. coeruleus* cells we used were lab-grown, the *S. stipatus* spec. nov. cells were wild-caught, and this difference in rearing might influence habituation abilities. Future experiments using wild-caught cells of both species can confirm our observed habituation patterns. Additionally, we have yet to explore whether *S. stipatus* spec. nov. shares the regenerative ability of *S. coeruleus* and several other *Stentor* species.

Our characterization of *S. stipatus* spec. nov. adds to our understanding of common principles of complex behavior that are present in single-celled organisms. Furthermore, our morphological and environmental descriptions contribute to our knowledge of protist natural history, which remains relatively understudied. Finally, since ciliate biomass is a major component of aquatic ecosystems, our exploration of ciliate diversity helps us understand the roles of complex cellular behavior in carving out ecological niches.

## CONCLUSIONS

In the present paper we describe a heterotrich ciliate stentor, *Stentor stipatus* spec. nov., collected from the iron-rich waters of the white cedar swamp in the Two Ponds Conservation Area of Falmouth, Massachusetts. Despite resemblance to *S. amethystinus*, we provide morphological and phylogenetic evidence that *S. stipatus* spec. nov. is likely a novel species. We further demonstrate that this species of *Stentor* can habituate, albeit faster than the archetypal species, *S. coeruleus*, and exhibits strong positive phototaxis, like *S. pyriformis*, which fluctuates in a consistently diurnal manner. Our work establishes *S. stipatus* spec. nov. as a new species of interest for studies in cell behavior and circadian regulation.

## DECLARATIONS

## Ethics approval and consent to participate

Not applicable.

## Consent for publication

Not applicable.

## Data availability

Image and video files analyzed for phototaxis and habituation experiments are available upon request. 18S SSU DNA sequences are included in the published work and are available on GenBank with the following accession numbers: Sequence 1 (*S. coeruleus*; wild strain) PX056129, Sequence 2 (*S. muelleri*; wild strain) PX056130, Sequence 3 (*S. pyriformis*; wild strain) PX056131, Sequence 4 (*S. stipatus*; spec. nov., wild strain) PX056132, Sequence 5 (*S. stipatus*; spec. nov., wild strain) PX056133, Sequence 6 (*S. stipatus*; spec. nov., wild strain) PX056134, Sequence 7 (*S. stipatus* spec. nov. wild strain) PX393983. *S. stipatus* spec. nov. as described in this work, has been registered on ZooBank, LSID: urn:lsid:zoobank.org:pub:0C20B702-EE37-47C9-8DDE-417C2B39CBC4.

## Author’s contributions

DHR performed the habituation experiments as well as the phylogenetic analysis and drafted the manuscript. AA, ET, AM, CV and BL all contributed to single cell 18S SSU sequencing. BL also contributed to phylogenetic tree assembly. WFM supervised DHR and provided support and guidance for experimental design as well as feedback and edits on the manuscript. KR performed phototaxis and action spectrum experiments. DBC performed phototaxis and action spectrum experiments, did the imaging for morphological analysis, supervised KR, and drafted and edited the manuscript. All authors read and approved of the final manuscript.

## Acknowledgements

Initial discovery and collection of this novel *Stentor* species was made possible through funding by the Grass Foundation in 2023. Initial culturing and most of the phototaxis experiments were performed at the Marine Biological Laboratory (MBL) in the summers of 2023 and 2024. We also thank Dr. Mark Slabodnick and Dr. Anton Suvorov for guidance on phylogenetic analysis.

## ADDITIONAL FILES

Additional file 1: **Movie 1**. Cortical enrichment of autofluorescent algae in *S. stipatus* spec. nov. (.mp4)

Additional file 2: **Movie 2**. Random movement of *S. stipatus* spec. nov. in uniform white light. (.mp4)

Additional file 3: **Movie 3**. Directed movement of *S. stipatus* spec. nov. in asymmetric white light. (.mp4)

Additional file 4: **Movie 4**. Distribution of cortical algae in *S. stipatus* spec. nov. after centrifugation. (.mp4)

Additional file 5: **Movie 5**. Abnormal swimming of centrifuged *S. stipatus* spec. nov. in asymmetric light. (.mp4)

Additional file 6: **Movie 6**. Redistribution of pigmentation after recovery from centrifugation. (.mp4)

Additional file 7: **Movie 7**. Normal swimming of *S. stipatus* spec. nov. in asymmetric light following recovery. (.mp4)

Additional file 8: **File S1**. Sequence alignment generated by MAFFT. (.fasta) Additional file 9: File S2. 18S sequences for *Stentor* species. (.fasta)

Additional file 10: Files S3. Treefile of aligned 18S sequences generated by IQTREE(v2.0.7)

